# Estimates of genetic load suggest frequent purging of deleterious alleles in small populations

**DOI:** 10.1101/696831

**Authors:** Tom van der Valk, Marc de Manuel, Tomas Marques-Bonet, Katerina Guschanski

## Abstract

It is commonly thought that declining populations will experience negative genetic consequences as a result of increased inbreeding. Here we estimated the average deleteriousness of derived alleles in a range of mammals and found that species with historically small population size and low genetic diversity often have lower genetic load than species with large population sizes. This is likely the result of genetic purging – the more efficient removal of partially deleterious recessive alleles from inbred populations. Our findings suggest that genetic purging occurs over long evolutionary time frames, and therefore rapid population declines are likely to dis-proportionally increase mutational load in species with high diversity, as they carry many deleterious alleles that can reach fixation before genetic purging can remove them.

## Main

Small inbred populations of wild animals frequently show lower survival, less efficient mating and lower reproductive success than large outbred populations (1) as a consequence of high levels of genome-wide homozygosity at loci with partially recessive deleterious alleles (2). Such negative fitness consequences of inbreeding have been directly confirmed on the genomic level (3–6). Nevertheless, many animals have survived in small populations for thousands of generations without apparent strong negative fitness effects. A suggested explanation for this phenomenon is genetic purging — the increased efficiency of purifying selection at removing partially recessive deleterious alleles in inbred populations (7). Whereas in large populations partially recessive deleterious alleles are mostly found at low frequency, these alleles can drift to high frequency in small populations (2). Mating between related individuals subsequently brings recessive alleles in a homozygous state, exposing them to purifying selection and thus leading to their more efficient removal over time (2). Although genetic purging has been shown to act in several animal populations (8–11), it remains largely unexplored to what extent this process represents a prominent evolutionary force. Wild animal populations across the globe experience rapid declines (12, 13), which are often followed by increased rates of inbreeding and the resulting genetic consequences can directly contribute to their extinction (14). Understanding under what circumstances genetic purging acts and how common it is among endangered populations could therefore help to identify species facing the most severe genetic consequences of population declines and prioritize them for conservation interventions.

Here, we used genomic data to estimate the prevalence of genetic purging in wild mammalian populations, as mammals are among the most affected by human-induced population declines (12). By calculating phylogenetic sequence conservation we aimed to identify genomic sites under strong evolutionary constraints, as mutations at these sites are expected to bear negative fitness consequences (15). Genomic sites that remained conserved during millions of years of evolution are expected to be functionally important, and therefore mutations at such sites can serve as a proxy for genetic load – the reduction of population mean fitness due to genetic factors (16, 17). Using a panel of 100 mammalian reference genomes, comprising all major mammalian lineages, we calculated the genomic evolutionary rate profiling (GERP) scores as the number of rejected substitutions, i.e. substitutions that would have occurred if the focal genomic element was neutral but did not occur because it has been under selective constraints (18). Mutations at highly conserved genomic sites (those sites with high GERP-scores) are likely deleterious, whereas those at low GERP-scores are expected to be mostly neutral. We then used these GERP-scores to estimate individual relative genetic load in 655 individuals belonging to 41 different mammalian species, using publicly available whole genome re-sequencing data.

## Results and Discussion

### GERP scores indicate conserved genomic regions

We used a short-read mapping-based approach for the GERP-score calculations (see methods, SI Appendix, Fig. S1, S2) and validated its accuracy using five independent approaches. First, for the human genome, we obtained a high correlation between the GERP-scores previously calculated from a 44 whole-genome vertebrate alignment (18) and those obtained with our pipeline (Pearson correlation *r* = 0.944) (SI Appendix, Fig. S3). Second, our calculated GERP-scores are four to six times higher within exonic regions, known to be highly conserved and under purifying selection in vertebrates (19), than in intronic regions (P < 2.2 · 10^−16^) (SI Appendix, Fig. S4). Third, the majority of within-population variable sites (88% ±SE 5% across all species) are found at the lowest 10% of GERP-scores, suggesting that low GERP-scores mostly reflect neutrally evolving regions that are segregating within populations, whereas polymorphisms at high GERP-scores are mostly removed from the populations (SI Appendix, Fig. S5). Fourth, we observe that derived alleles at high GERP-scores are more often found in the heterozygous state, whereas at low GERP-scores they often appear in the homozygous state (SI Appendix, Fig. S6), suggesting that many derived alleles at high GERP-scores present in a population are likely to be recessive deleterious. Finally, GERP scores are on average higher at amino-acid changing sites that are expected to have a deleterious impact on the protein, as calculated with SIFT (20), compared to more tolerated amino-acid changing positions (SI Appendix, Fig. S7), providing an independent support for the likely negative effects of mutations in conserved parts of the genome. Taken together, these results are consistent with the expected dynamics of partially deleterious recessive alleles (21).

### Population size and levels of inbreeding correlate with genetic load

Within species differences in genetic load are often estimated as the total number of putative deleterious alleles per genome (11, 17). However, when comparing different species, obtaining unbiased estimates of genetic load that are based on sums becomes challenging, as species differ from each other in genome size, mutation rate and generation time and these factors lead to differences in the number of variable sites independent of genetic load. Thus, to allow comparisons across species, we calculated individual genetic load as the average GERP score of all derived alleles (Methods, Figure 1A). This measure is independent of the number of derived alleles identified in the genome, but rather reflects the distribution of conservation scores of the derived alleles. The rationale behind this approach is the following: As purifying selection acts on deleterious alleles, here identified as those at highly conserved sites, the distribution of derived alleles shifts towards lower GERP scores. The stronger the purifying selection, the stronger the shift of the distribution towards lower GERP scores. Thus the average GERP score of the derived alleles in a genome reflects the strength of purifying selection experienced by a given species over time. As we count all derived alleles in the genome, independent of zygosity, we implicitly assume co-dominance of these alleles (but see SI Appendix, Fig. S14 for different considerations of the dominance coefficient). Using this measure, we found that estimates of genetic load differ among the studied mammalian species, but are very similar for individuals belonging to the same species (Fig. 1A). The low inter-individual variation in genetic load (±SD 1.3%) suggests that this measure reflects long-term evolutionary processes (e.g. over hundreds of generations). We confirmed that our inferences are robust with respect to the outgroup species used to determine the ancestral state of alleles, as well as to the species used in the genome alignments for GERP-score calculations (SI Appendix, Fig. S8, S9, S10), suggesting that between-species differences are not driven by underlying technical artifacts.

**Fig. 1.**
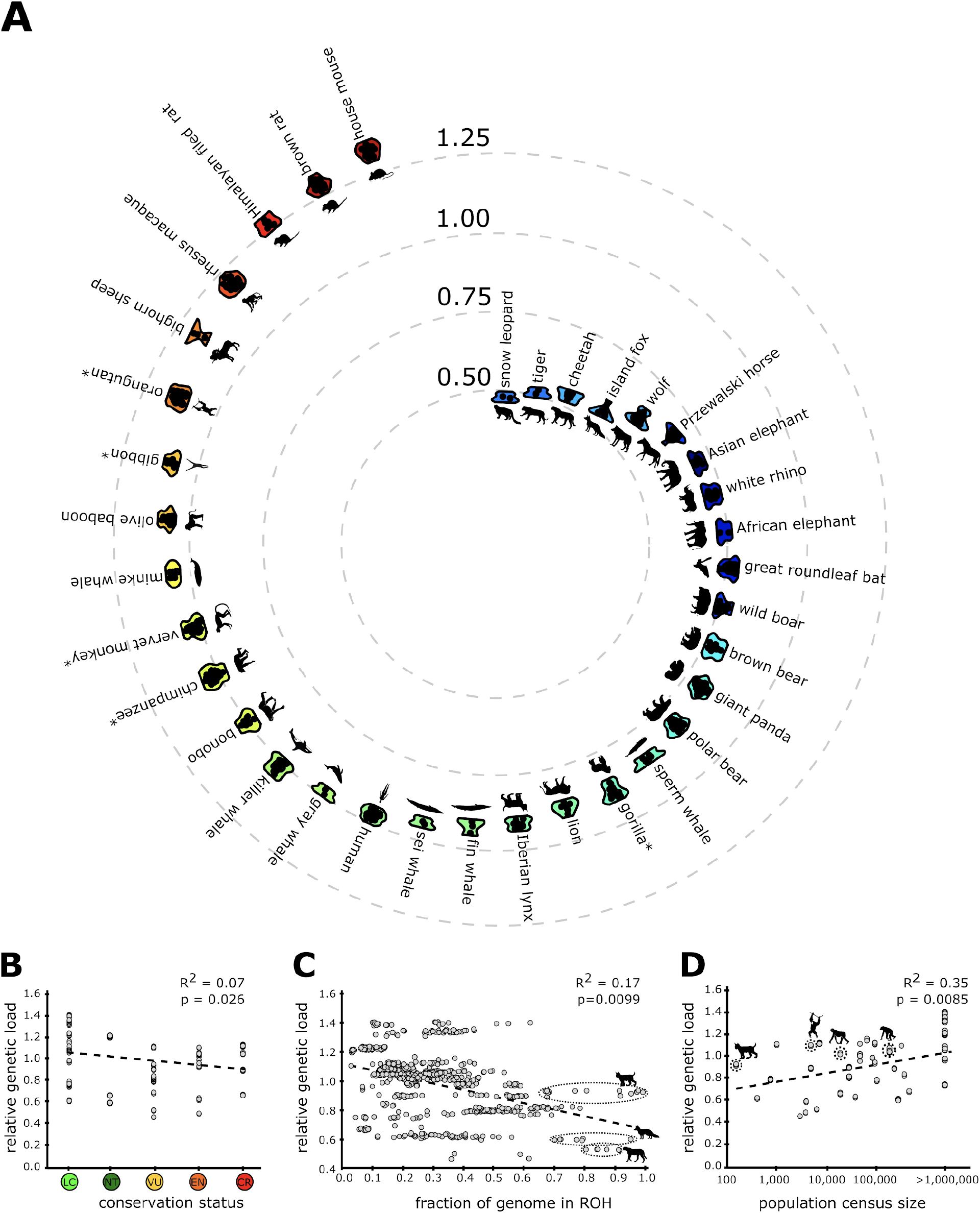
Relative genetic load in mammals. (**A**) Genetic load is depicted as the average GERP-score of the derived allele for each individual within a species. Several closely related species (Sumatran and Bornean orang-utans, gibbons, vervet monkeys, and eastern and western gorillas) are grouped together for clarity (marked by the asterisks). (**B**) Relative genetic load is not well explained by conservation status. LC: least concern, NT: near threatened, VU: vulnerable, EN: endangered, CR: critically endangered. **(C)** Relative genetic load is negatively correlated with inbreeding. Species with recurrent bottlenecks and/or small population size show low load despite high inbreeding (e.g. cheetah and island fox). In contrast, some highly inbred species, which experienced recent dramatic population decline show disproportionally high genetic load (e.g. Iberian lynx). **(D)** Genetic load is generally higher in species with large census population size (species with population size above 1 million are grouped together for clarity). However, some species with historically large population sizes and recent strong declines (e.g. Sumatran orangutan, Iberian lynx, chimpanzee, bonobo) show elevated genetic load given their population size. Each grey dot represents a species in (B) and (D) and an individual genome in (C). Dotted lines depict the best fitting linear intercept.

We sought to explore whether measures of genetic load can be meaningfully applied in species conservation. Endangered species often have small population sizes, low genetic diversity and high inbreeding (14) and as a result are expected to carry high genetic load (1). Although we find that genetic load and species conservation status correlate, the relationship is weak (R=0.07, p=0.026, Fig. 1B) and goes in the opposite direction of the above expectation, with lower genetic load in species of greater conservation concern.

As conservation status does not correlate with genetic diversity and only weakly with population size (22), we investigated whether these measures may be explained by differences in genetic load among species. We observed a weak inverse relationship between genetic load and inbreeding (R=0.17, p=0.0099), calculated as proportion of the genome in runs of homozygosity (Fig. 1C). Many species with low genetic load, i.e. those with most derived alleles at only weakly conserved sites, have high proportions of their genome in runs of homozygosity (e.g. snow leopard, tiger, island fox, wolves, cheetah, Fig. 1C, SI Appendix, Dataset S1). Conversely, species with high genetic load frequently have a low genome-wide rate of homozygosity (e.g. house mouse, brown rat, Himalayan rat, vervet monkey, olive baboon, rhesus macaque, SI Appendix, Dataset S1). However, because in contrast to intra-specific measures of genetic load, the level of inbreeding can change rapidly within only a few generations, which is exemplified by the high degree of intra-specific variation in inbreeding (±SD 27%) (SI Appendix, Dataset S1), the correlation between genetic load and inbreeding is only moderate (Fig. 1C).

The ratio of genetic load calculated for genomic regions outside and within runs of homozygosity also differed across species (SI Appendix, Fig. S11). It was smaller in large outbred populations, signifying higher genetic load within runs of homozygosity than outside (e.g. houses mouse, vervet monkey), and larger in populations with a long history of inbreeding and long-term small population sizes, showing lower genetic load within runs of homozygosity than outside (e.g. snow leopard) (SI Appendix, Fig. S11). Overall, there was a significant correlation between this ratio and total genetic load in the species, such that species with higher ratios showed lower genetic load (SI Appendix, Fig. S12). This observation is consistent with the action of purging, where partially recessive deleterious alleles are removed from the population through long-term inbreeding and exposure to purifying selection, since within populations certain genomic regions are more frequently found in homozygosity than others (23). In contrast, partially deleterious alleles persist in large populations as segregation load, as they rarely and inconsistently appear in a homozygous state in any given individual.

Contrary to the notion that small populations have high genetic load (24), we observe a positive relationship between genetic load and population size (Fig. 1D, SI Appendix, Dataset S1, R=0.35, p=0.0085). Generally, species with small population size have lower genetic load than species with large population sizes (Fig. 1D, SI Appendix, Dataset S1), suggesting that purging of deleterious alleles can be an important evolutionary force. This relationship was only slightly affected by the dominance coefficient of the derived alleles, with larger populations showing higher genetic load if the derived alleles were considered dominant (SI Appendix, Fig. S13). In small populations, genetic load was almost independent of the dominance coefficient, as most derived alleles are in a homozygous state and thus exposed to selection. Despite generally lower load in small populations, we observe relatively high genetic load in several species with historically large population sizes that have experienced dramatic recent population declines (e.g. chimpanzees, orangutans, bonobos and Iberian lynx, Fig. 1D) (25). This corroborates recent findings from genetic simulations, which demonstrate that strong declines in population size disproportionally affect ancestrally large populations (26). Finally, after controlling for the phylogenetic relationships among the studied species (27), the positive correlation between population size and genetic load remains but is less pronounced (R=0.153, p = 0.023, SI Appendix, Fig. S14). This is consistent with our earlier observation that changes in genetic load seem to occur over long evolutionary time-frames.

### Fixation of deleterious alleles

As selection can only act on variation, deleterious alleles that are fixed within a population are especially problematic for long-term population viability. We thus estimated the fraction of fixed derived alleles stratified by GERP-score for all species with at least five individuals in our dataset (Fig. 2a). Generally, we find that species with low genetic load carry few derived alleles at high GERP-scores (e.g. cheetah, island fox, Przewalski horse) (Fig. 2b), however, these alleles frequently appear to be fixed in the population (Fig. 2a). In contrast, although some populations with high genetic load (e.g. house mouse, brown rat, Himalayan field rat, European rabbit, vervet monkey, olive baboon, rhesus macaque) carry a high proportion of putatively deleterious alleles at high GERP scores (Fig. 2b), the majority of these are at low frequency and less likely to appear in the homozygous state in any given individual (Fig. 2a). Thus, while purging removes strongly deleterious alleles in highly inbred species, some deleterious alleles nonetheless reach fixation, which can subsequently lead to negative fitness consequences without the opportunity for additional genetic purging. This could also explain why inbreeding depression has been reported in the cheetah and (Swedish) wolves despite the relatively low overall genetic load (28, 29). Taken together, these observations are especially worrying for genetically diverse populations that experience rapid population declines, as we show that in species with high genetic diversity more derived alleles are found at high GERP scores and thus a high proportion of these putatively deleterious alleles could reach fixation before genetic purging can act (see also (24)).

**Fig. 2.**
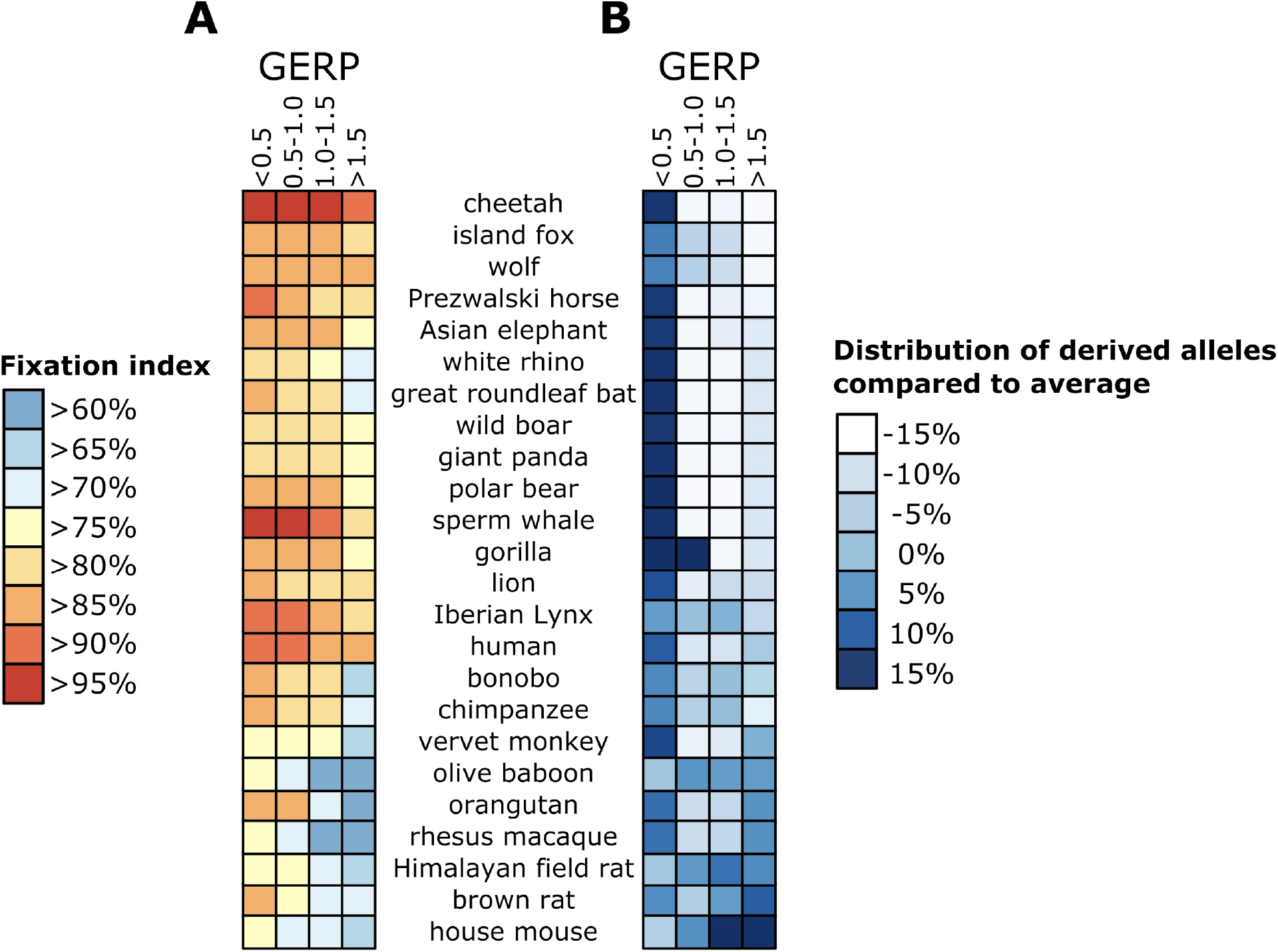
Fixation of derived alleles. Species are ranked by average GERP-score of the derived alleles (following Fig. 1A) from the lowest (at the top of the graph) to highest (bottom of the graph). (A) Percentage of derived alleles that are fixed within the species for a given GERP-score bin. E.g. most derived alleles in cheetah are fixed, even at high GERP-score, whereas in house mouse many derived alleles are segregating in the population. (B) Distribution of derived alleles in each species for each GERP-score bin compared to the average over all species. Darker colors represent relatively more derived alleles within the given GERP-score class. E.g. cheetahs have more derived alleles at low GERP-scores, whereas derived alleles in the house mouse are more frequent at high GERP-scores.

### Genetic implications for conservation of small populations

Our study describes a general trend of increasing genetic load with population size and outbreeding (Fig. 1c-d). This pattern allows the identification of populations and species that could represent potential targets for conservation interventions as those falling above the trendline and thus showing higher genetic load than would be expected given their level of inbreeding or their population size. For instance, individuals of the Iberian lynx show elevated genetic load given their level of inbreeding, which is in accordance with their history of strong recent bottlenecks. Similarly, multiple great ape species (orangutans, bonobos, chimpanzees) show higher genetic load than would be expected given their current population size, which is in line with their rapid population declines. These species and populations could thus represent priorities for conservation interventions, as they appear to have a larger genetic load than other species of comparable population size and might therefore be vulnerable to inbreeding depression.

Deliberate purging by inducing inbreeding has previously been suggested as a management strategy for reducing inbreeding depression (30). Current evidence for its efficiency is limited (21), but some wild populations show evidence of successful purging. For instance, the Steward Island robins, birds with a long-term small population size, show no correlation between inbreeding and juvenile survival (31), whereas juvenile survival in the closely related Chatman Island robins, which went through a drastic recent bottleneck, is negatively associated with levels of inbreeding (31). The long-term small population size of the Stewart Island robin thus seems to have facilitated extensive purging, making this species more “adapted” to small population size. Although this shows that deliberate purging could potentially result in a positive outcome, several species with recent sharp population declines show little signs of genetic purging (e.g. orangutan, chimpanzee, lions). Therefore purging seems to be of limited efficacy during rapid population declines (and the consequently sharp rise in inbreeding). Simulations also suggest that although inbreeding can be effective in purging lethal alleles within a few generations, the removal of partially recessive deleterious alleles with smaller fitness effects requires long-lasting and slow inbreeding (8, 32). As such, deliberate purging seems to be only an effective strategy for short-term conservation if the aim is to remove (nearly) lethal alleles from a population (33). In addition, experimental studies show that rapid serial bottlenecks can result in a highly increased risk of extinction (34). Given all this evidence, it is conceivable that small inbred populations that are alive today represent those species that have effectively purged deleterious alleles, whereas rapid inbreeding has led to the extinction of many other populations. Finally, deliberate inbreeding will result in loss of genetic diversity and thus limit future adaptive potential of populations (35, 36). In management of captive breeding and restoration programmes, deliberate inbreeding is thus a risky strategy with likely negative outcomes in most cases.

The higher genetic load of individuals from large populations represents an important aspect to be considered for the commonly employed conservation strategy of genetic rescue. The main objective of genetic rescue is to mitigate inbreeding depression by introducing genetic diversity and beneficial variants through translocation of outbred individuals into an inbred population. Although genetic rescue has been shown to increase population fitness in the short-term for a range of mammals (37, 38), the long-term effects can be negative. This is exemplified by the collapse of the Isle Royale wolves, a population that maintained good population viability for decades. After interbreeding with a mainland wolf migrant, the Isle Royale wolves initially showed higher reproductive success. However, subsequent inbreeding in this population eventually resulted in the increase in frequency of deleterious alleles, introduced by the immigrant, and eventual marked decline of the population (5). A similar event occurred in the inbred Scandinavian wolf population following the arrival of a single immigrant wolf, but subsequent additional immigration has resulted in additional outbreeding and a period of recent population growth (39). Recurrent introductions of genetically diverse individuals into the population was also proposed as an explanation as to why domestic chickens, in contrast to most domesticated species, do not shower higher genetic load than their wild counterparts (40). Our analyses demonstrate that individuals from large outbred populations often carry higher genetic load, present as partially recessive segregating deleterious alleles. The introduction of an outbred individual with an array of deleterious alleles into an inbred population that has experienced genetic purging may thus have negative consequences. These consequences can possibly be avoided by selecting individuals with low genetic load to be used for genetic rescue (26). However, if not followed by a clearly delineated long-term plan to reduce inbreeding, for instance through repeated introductions, the addition of strongly deleterious alleles into a small population with low capacity for long-term purifying selection and purging may lead to fixation of these alleles with detrimental effects for population survival.

## Methods

### Single nucleotide variant calling

We obtained published re-sequencing data for 655 mammalian genomes from 41 species and mapped these to the phylogenetically closest available reference genome for each species (SI Appendix, Dataset S1) using bwa mem v0.7.17 (41). In total, 27 reference genomes were used for this task (SI Appendix, Dataset S1). We then obtained and filtered variant calls for each individual using GATK HaplotypeCaller v3.8 following the “short variant discovery best-practices guidelines” including “hard filtering” (42). Additionally, we only kept within-species bi-allelic sites and removed all indels and sites below one third and above three times the genome-wide autosomal coverage (43).

### Genomic Evolutionary Rate Profiling

We used the software GERP++ (Genomic Evolutionary Rate Profiling) to calculate the number of “rejected substitutions” (a proxy for evolutionary constraints) for each site in the same 27 reference genomes that were used in mapping of the re-sequencing data (SI Appendix, Dataset S1, SI Appendix, Fig. S1) (18). GERP++ estimates the number of substitutions that would have occurred if the site was neutral given a multi-species sequence alignment and the divergence time estimates between the aligned species as provided in (44). A GERP-score, the number of rejected substitutions at a genomic site, is thus a measure of constraint that reflects the strength of past purifying selection at a particular locus. To calculate GERP-scores for a given focal reference genome, we used 100 non-domesticated mammalian de-novo assembled genomes (SI Appendix, Dataset S2, SI Appendix, Fig. S2), as domesticated species might give a biased estimate of purifying selection. Each individual genome sequence was converted into short FASTQ reads by sliding across the genome in non-overlapping windows of 50 base pairs and transforming each window into a separate FASTQ read. The resulting FASTQ reads from the 100 mammalian genomes were then mapped to each respective focal reference genome with bwa mem v0.7.17, slightly lowering the mismatch penalty (-B 3) and removing reads that mapped to multiple regions. Mapped reads were realigned around indels using GATK IndelRealigner (45, 46). Next, we converted the mapped reads into a haploid FASTA consensus sequence (i.e. 100 times for each reference genome), excluding all sites with depth above one (as such sites contain at least one mismapped read). GERP++ was then used to calculate the number of rejected substitutions at all sites in the reference using the concatenated FASTA files and the species divergence time estimates from (44) (SI Appendix, Fig. S2), excluding the focal reference from the calculation. Missing bases within the concatenated alignment were treated as non-conserved (i.e. sites for which only few reads mapped obtain low GERP scores). We excluded all sites for which the focal reference FASTQ reads did not map to themselves and sites with negative GERP-scores (as these most likely represent errors) and subsequently scaled all scores to a range from 0 to 2. Sites that are identical between species and have thus been preserved over long evolutionary time result in high GERP-scores (SI Appendix, Fig. S1). Thus, high GERP-scores are only obtained for regions, where the majority of the 99 mammalian genomes (100 minus the focal reference) map to the respective reference.

### Effects of phylogeny on estimated GERP scores

To evaluate whether GERP score estimates could be affected by the availability or lack of sequenced genomes from closely related species (e.g. primates with many available genomic sequences from close relatives versus Proboscidea for which most of the sequenced species are evolutionary distant), we simulated a tree with the same number of species (N=40) but different evolutionary distances between the focal species and all other species compared (SI Appendix, Fig. S10). We calculated the GERP score for the human genome for three phylogenetic trees that differed in the average evolutionary distance between the focal genome (human) and all other species. Mean evolutionary distance ranged from 33.6 million years (SI Appendix, Fig. S10 Tree 1 containing only primates and two outgroups) to 57.3 million years (SI Appendix, Fig. S10 Tree 2) and 72 million years (SI Appendix, Fig. S10 Tree 3). We observed a close similarity and strong correlation between estimated GERP scores independent of the evolutionary distance separating the focal species from other species in the phylogenetic tree, suggesting that any biases arising during the calculation of GERP scores between densely and sparsely represented parts of the mammalian tree of life are likely to be minimal.

### Ancestral allele inference

We called the ancestral allele at each site as the variant present in the phylogenetically closest outgroup. The derived alleles in each individual from the study dataset was then inferred against the called ancestral allele. By using only one outgroup we retained the highest number of sites to be analysed (the more outgroups are added, the fewer sites will be mapped across all outgroups). We estimated the effect of using one or multiple outgroups for the ancestral allele inference by calling the majority allele among the mapped reads for 1, 2 and 3 outgroups (a random base was choses if the allele frequency was equal) and show that this does not significantly change the estimates of genetic load (SI Appendix, Fig. S8, S9), as genomic sites with high GERP-scores are generally conserved and thus identical among all outgroup species (SI Appendix, Fig. S8).

### Relative genetic load

We estimated relative genetic load for each of the genomes as the average GERP-score of all derived alleles:

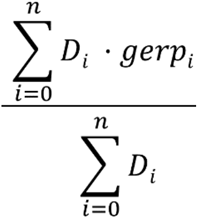

Where D_i_ represents the i^th^ derived allele and gerp_i_ the GERP-score for the i^th^ allele. We calculated averages to allow comparisons across species that differ in genome size and genome quality, as both these factors will affect the more frequently used genetic load estimates that sum across all deleterious mutations in the genome (11, 17). Whereas the score based on sums is highly appropriate for comparisons within a single species or across closely related species where variation in genome size and quality is negligible, the estimate based on means reflects the overall distribution of the mutations without the need to identify every single one of them. Under the assumption that new mutations occur randomly with respect to the genomic region, we expect that in species that experienced strong purifying selection, derived alleles are found mostly at non-conserved sites (low GERP-scores), whereas accumulation of deleterious variants should result in a higher fraction of derived alleles at high GERP-scores.

### Calculation of SIFT scores

SIFT-scores are a measure of the likelihood that an amino-acid change causes disruption of the protein function (20). We used SIFT 4G to identify all amino-acid changes classified as synonymous, deleterious and tolerated in the human, wolf and mouse genomes (as these have high quality genome annotation available) and calculated the distribution of GERP-scores among those three categories to test for correspondence between the two methods.

### Fixation of deleterious alleles

The fraction of fixed derived alleles was estimated for all species for which at least five individuals (e.g. 10 alleles) were present in our dataset. For species with more than five sequenced individuals, we randomly sampled 10 alleles at each site to exclude sample size bias. In both cases, we calculated the fraction of fixed derived alleles stratified by GERP-score.

### Individual inbreeding estimates

We used PLINK1.9 (47) to identify the fraction of the genome in runs of homozygosity longer than 100kb, a measure of inbreeding (F_ROH_), for all individuals with average genome coverage > 3X as in (48, 49). To this end, we ran sliding windows of 50 SNPs on the VCF files, requiring at least one SNP per 50kb. In each individual genome, we allowed for a maximum of one heterozygous and five missing calls per window before we considered the ROH to be broken. To account for differences in genome assembly qualities we restricted our analysis to contigs of at least 1 megabase.

### Phylogenetic independent contrast

We used the Phylogenetic independent contrast (PIC) (27) method from the R “ape” library to run a linear model on the correlations between population census size and genetic load, correcting for the fact that some species share a longer evolutionary history than others using the phylogenetic tree and species divergence times obtained from TimeTree (44)

## Acknowledgments

TvdV acknowledges support of a scholarship from the Foundation for Zoological Research. TMB is supported by funding from the European Research Council (ERC) under the European Union’s Horizon 2020 research and innovation programme (grant agreement No. 864203), BFU2017-86471-P (MINECO/FEDER, UE), “Unidad de Excelencia María de Maeztu”, funded by the AEI (CEX2018-000792-M), Howard Hughes International Early Career, Obra Social “La Caixa” and Secretaria d’Universitats i Recerca and CERCA Programme del Departament d’Economia i Coneixement de la Generalitat de Catalunya (GRC 2017 SGR 880). KG is supported by a Formas grant (2016-00835). We acknowledge support from the Uppsala Multidisciplinary Centre for Advanced Computational Science for access to the UPPMAX computational infrastructure.

## Author contributions

TvdV and MdM conceived the study and analysed the data. All authors interpreted the results. KG and TvdV wrote the manuscript with input from all authors.

## Data and materials availability

All data used in this study is available on the European Nucleotide archive (accession numbers are given in SI Appendix, Dataset S1). Scripts used in this study are available on github/tvdvalk.

## Supplementary Materials for

**Fig. S1.**
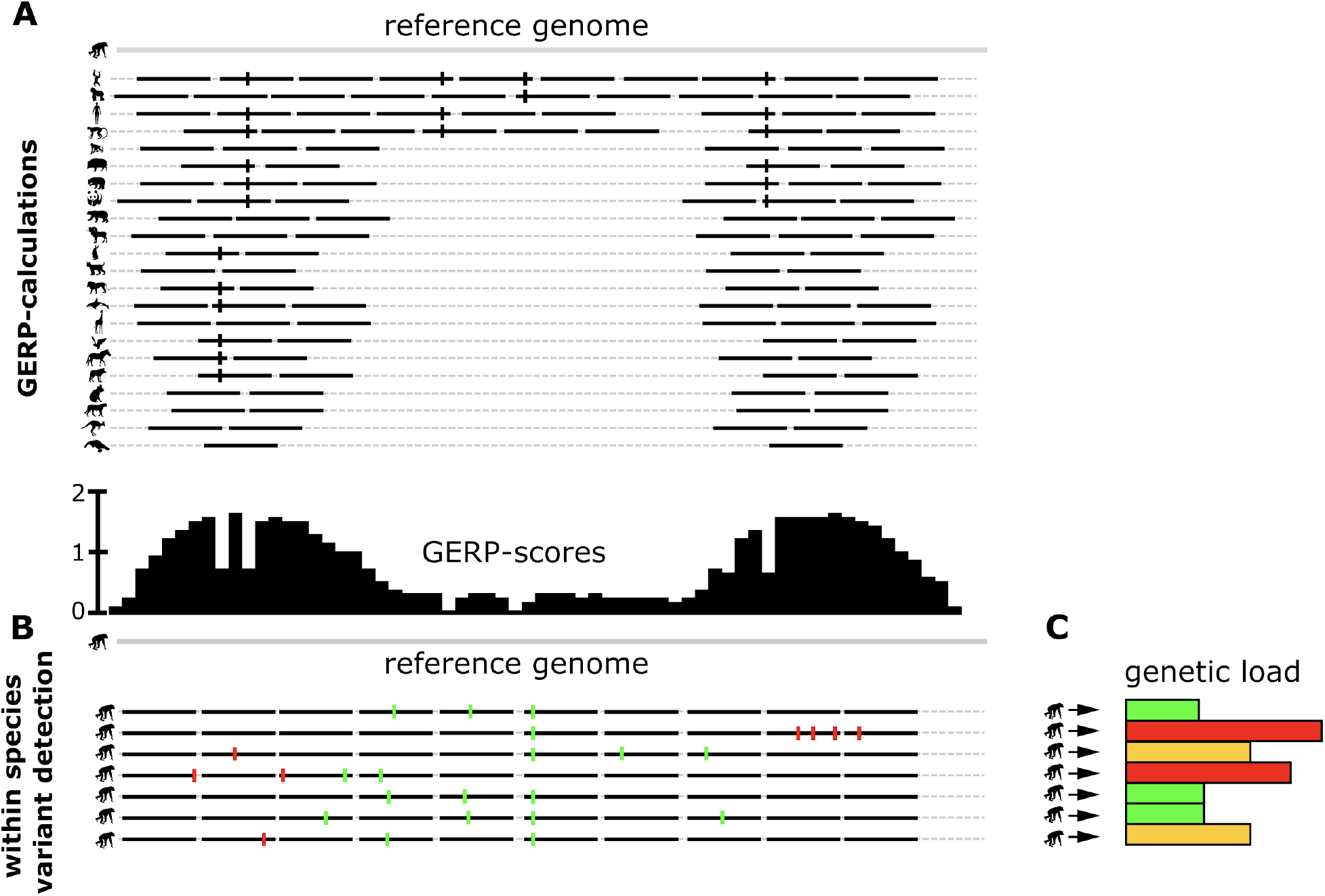
Schematic representation of the GERP-score pipeline. (A) A set of 100 de-novo assembled genomes is sliced into non-overlapping 50 base pair FASTQ read and aligned to the same reference as used for the within-species variance detection (SNP calling in (B)). A consensus sequence is then obtained for each of these 100 mapped genomes and GERP scores are subsequently calculated using the GERP++ software (excluding the focal reference from the calculation). Sites with few mapped reads or with a large proportion of variable alleles (depicted with vertical black bars on the individual reads) obtain low GERP scores, whereas sites identical among the majority of the mapped genomes obtain high GERP-scores. (B) Individual re-sequenced genomes from a population of a given study species are mapped to the reference genome (chimpanzee in this example) and SNPs are subsequently identified for each individual within the population following the GATK “short variant discovery best practise” guidelines. (C) The genetic load of the derived alleles identified in (B) can now be estimated. Derived alleles at highly conserved sites are more likely to have a negative fitness effect (depicted with the red vertical bars in B) compared to derived alleles at less conserved sites (green vertical bars in B). The average GERP-score of the derived alleles is used as a measure of the genetic load carried by each individual (red=high genetic load, orange=intermediate genetic load, green=low genetic load).

**Fig. S2.**
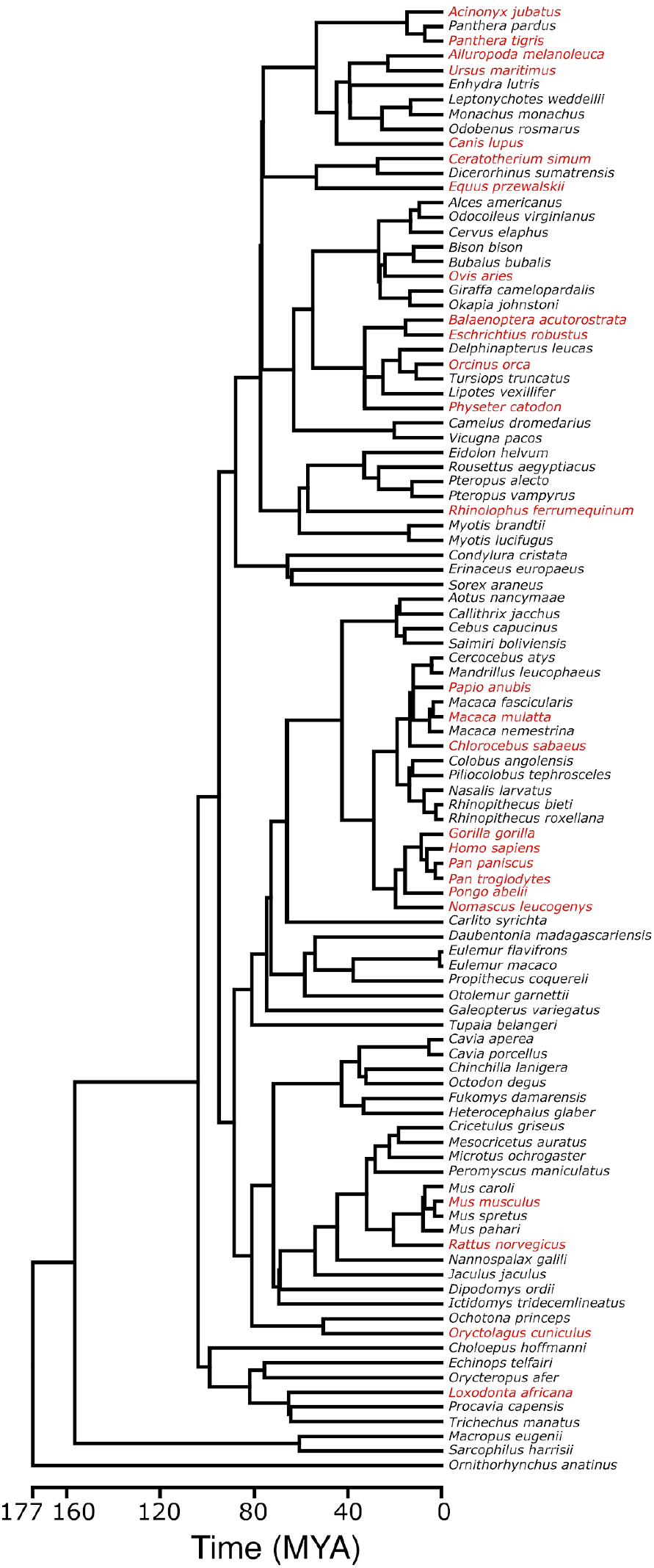
The 100 mammalian genomes and their divergence times estimates used for the GERP-score calculations. The divergence times between the species were obtained using the online software TimeTree, which gives a dated phylogeny from a list of species through automated literature searches (5). The genomes depicted in red were also used for the mapping of re-sequencing population-level data. Note, there are fewer such genomes than the number of studied species, as for species without a reference genome we used the closest available relative for mapping.

**Fig. S3.**
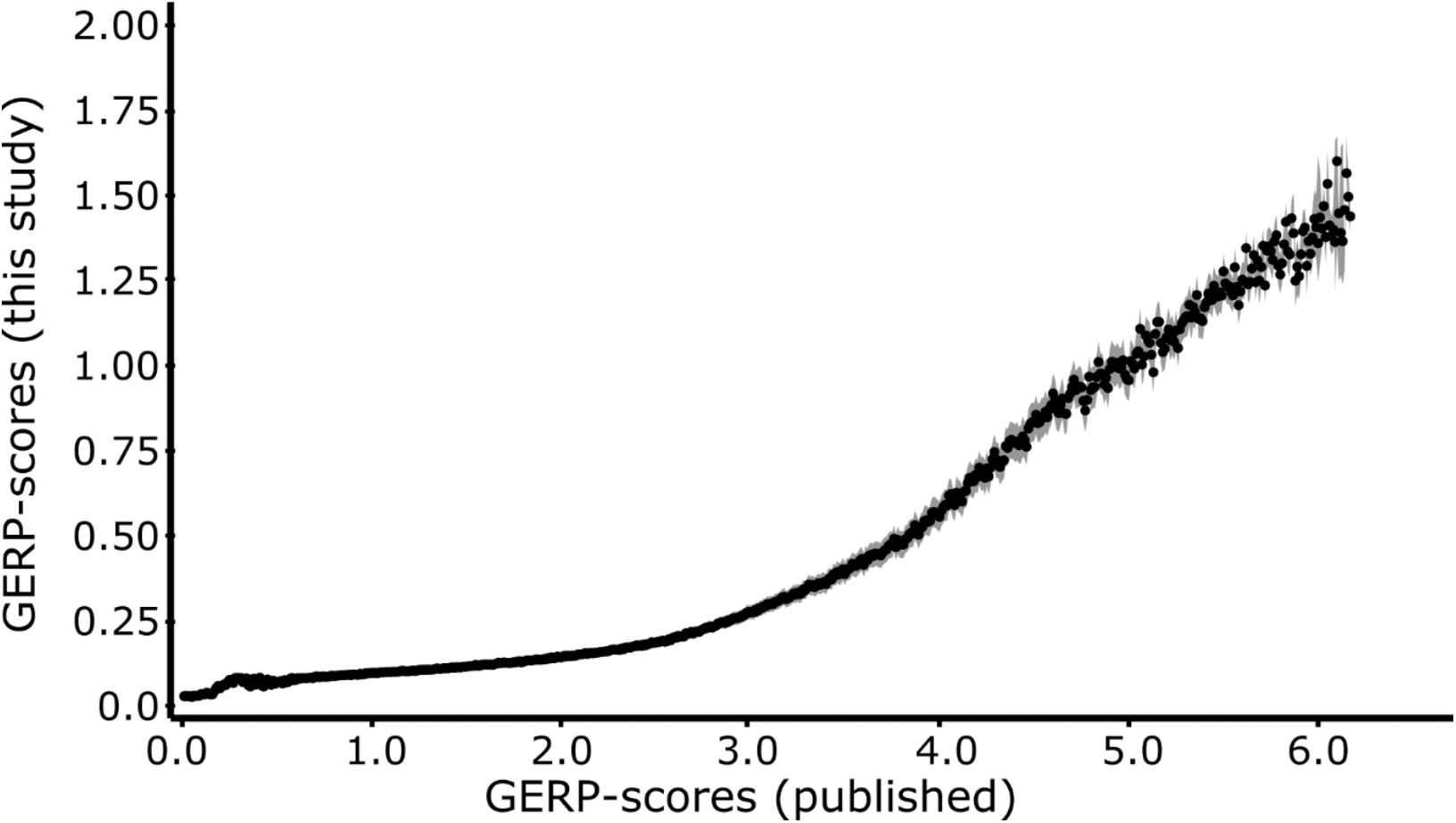
Correlation between previously published alignment-based GERP-scores and the GERP-scores calculated with the mapping-based approach in this study for the human genome (hg19). We binned all sites in the human genome by their published GERP-scores and calculated the average GERP-score for each bin of size 10 (black dots). Grey shaded area depicts ±1SD. Pearson correlation = 0.944, Spearman’s rank correlation = 0.997, p = P < 2.2 · 10^−16^. Note that we transformed our GERP-scores on a scale from 0 to 2, whereas the published scores are on the scale from 0 to 6.

**Fig. S4.**
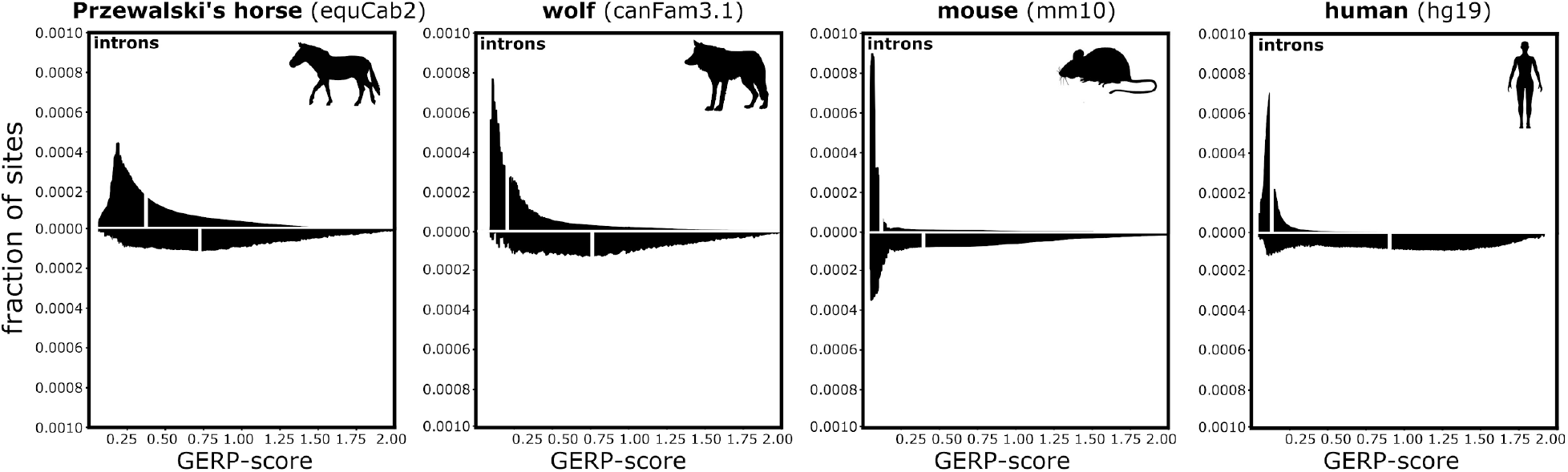
GERP-score of genomic partitions. The distribution of GERP-scores within introns (top) and exons (bottom) for 4 species with available high-quality reference genome annotations (used references between brackets). White lines within the plots depict the average GERP-score for a given genomic category. The highest GERP-scores are primarily found within exonic regions, with the average GERP-score in exons 4-6 times higher than within introns (P < 2.2 · 10^−16^ for all four species), which is consistent with stronger sequence conservation and selective constraints in exons compared on introns.

**Fig. S5.**
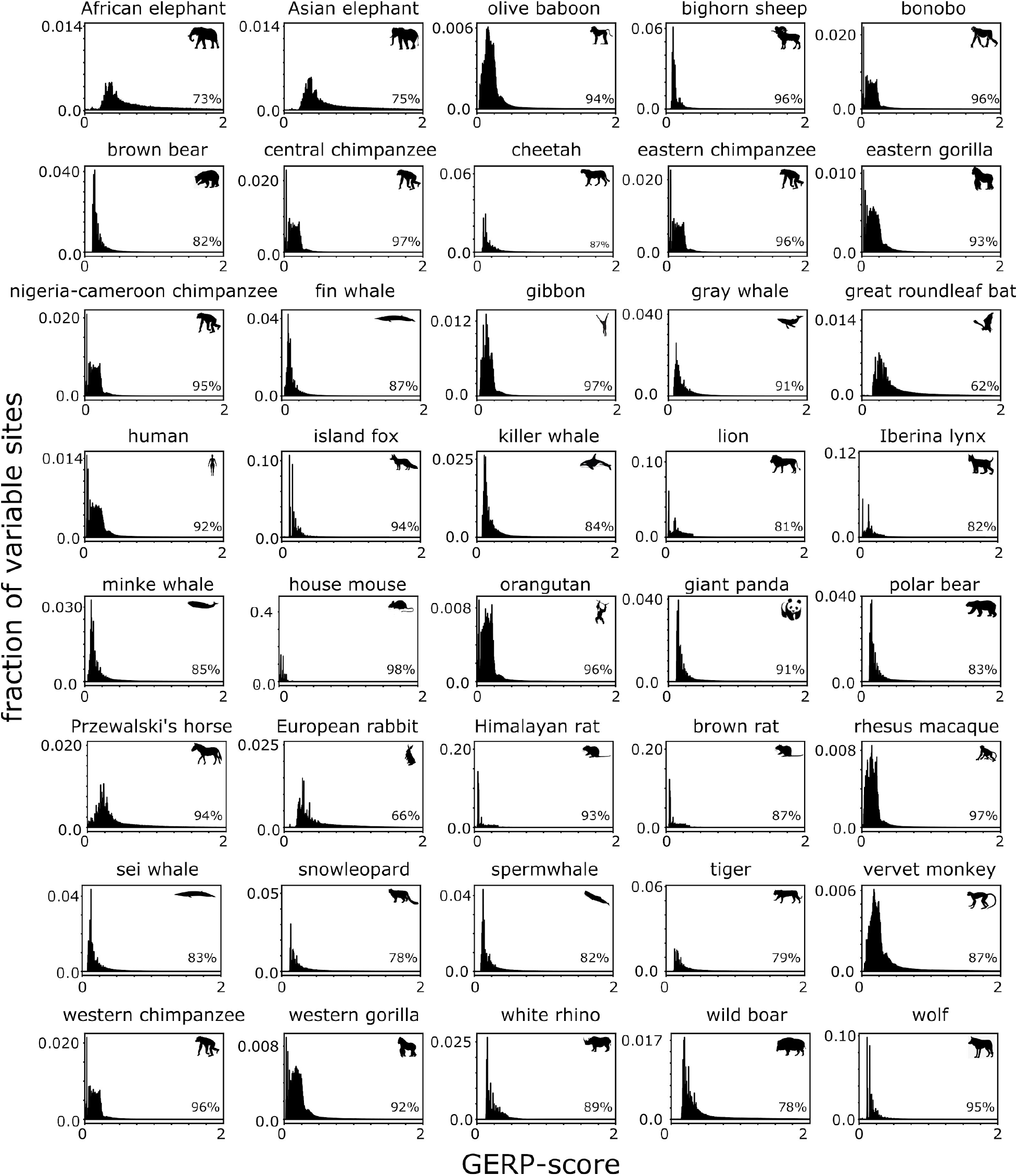
Distribution of segregating sites by GERP-score. The proportion of variable sites plotted against GERP scores, with the percentage found in the 10% lowest GERP-scores depicted in the bottom right corner. A large majority of variable sites are found at low GERP-scores and thus within fast evolving, likely neutral, regions.

**Fig. S6.**
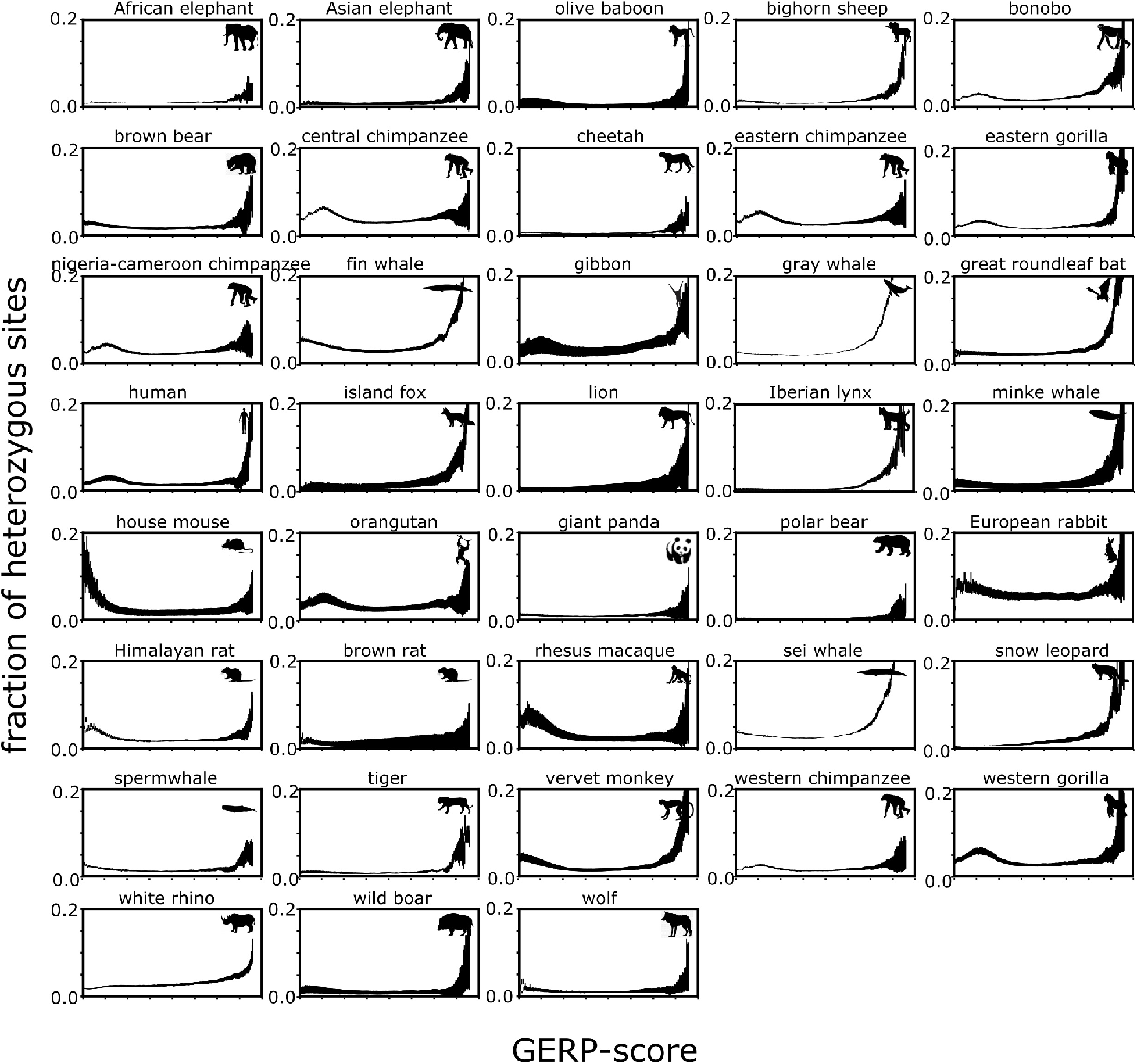
Proportion of heterozygous alleles out of total sites stratified by GERP-score. We included only samples with average genome wide coverage > 10X. X-axis is scaled from 0 to 2 for all species. The proportion of heterozygous derived alleles increases at high GERP-scores, consistent with them representing segregation load.

**Fig. S7.**
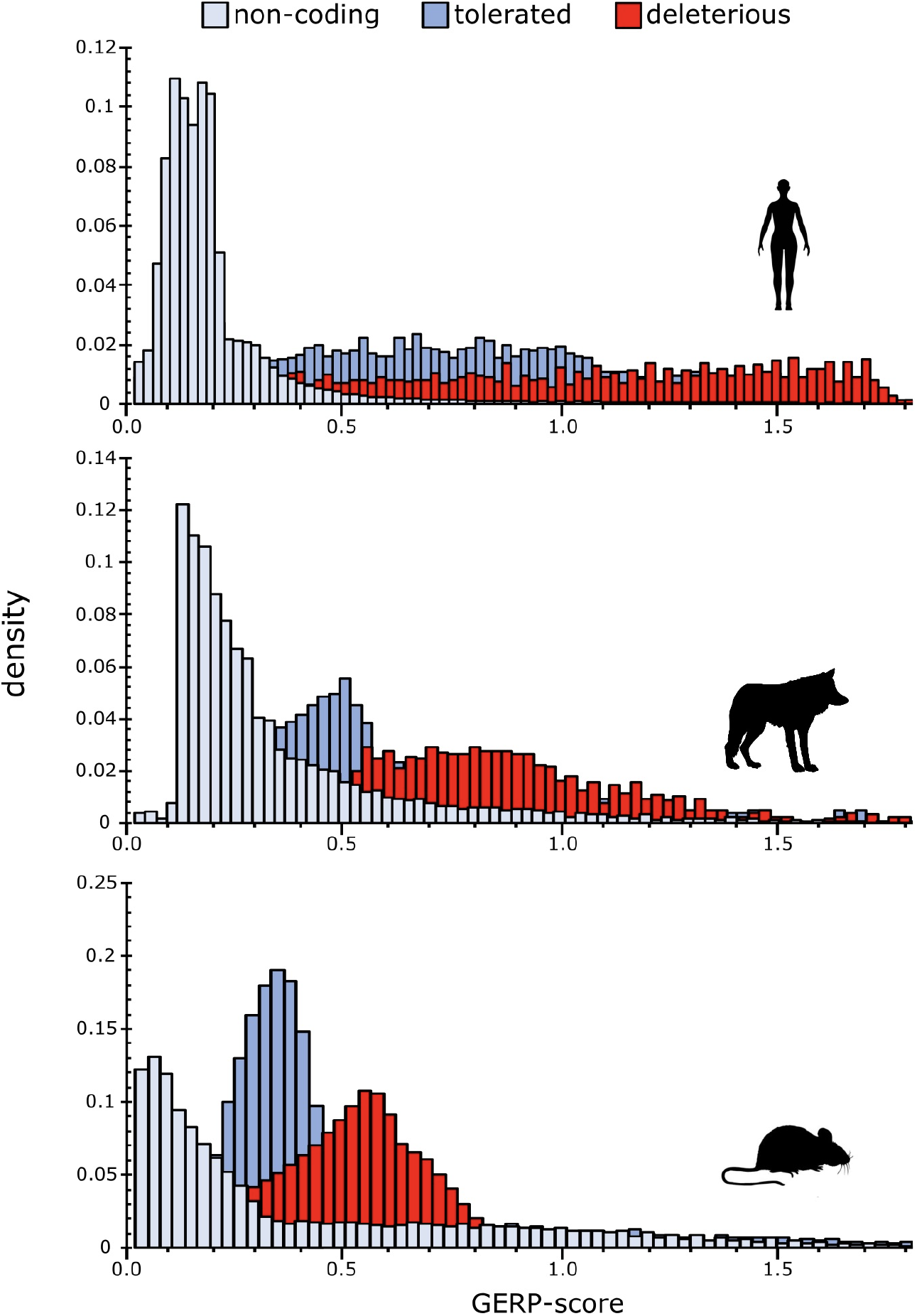
Correlation between SIFT and GERP-scores. Derived alleles in three phylogenetically distinct species with well-annotated genomes were classified with SIFT into non-coding, tolerated and deleterious variants. Generally, deleterious alleles are in genomic regions with higher GERP-scores than tolerated and non-coding alleles, providing independent confirmation for putatively deleterious effects of variants with high GERP-scores across mammals.

**Fig. S8.**
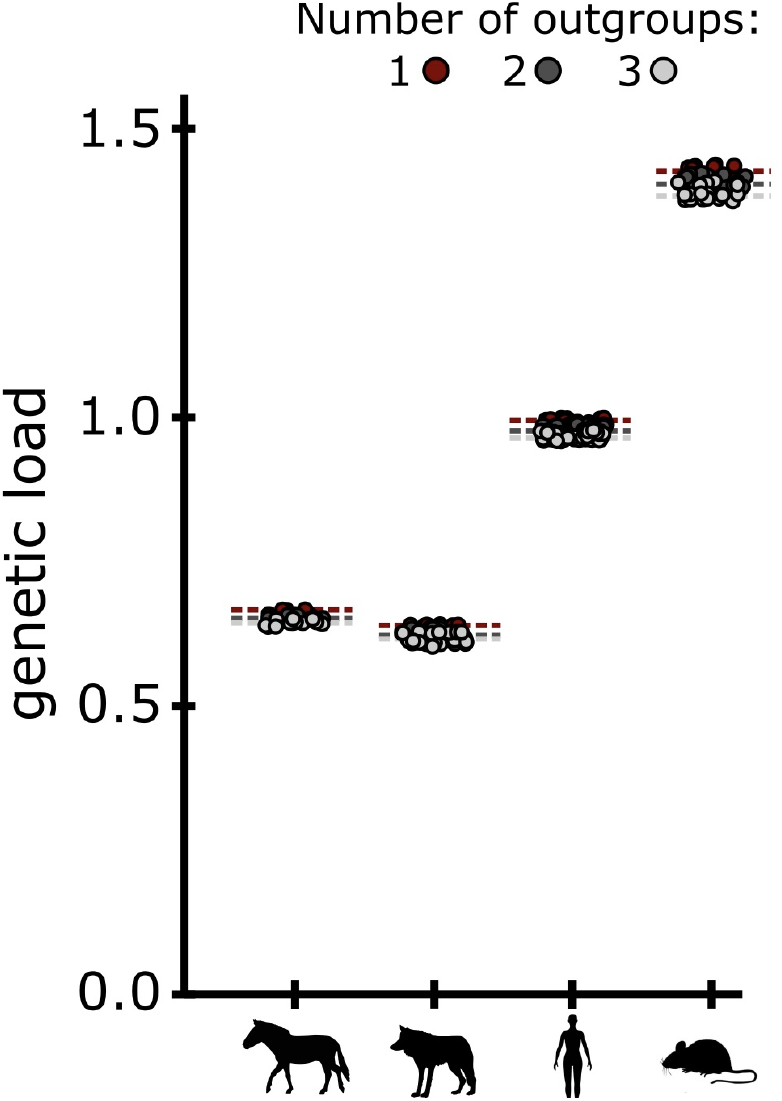
Relative genetic load calculated as the average GERP-score of the derived alleles using a different number of outgroups. We inferred the derived state by either using one outgroup or the majority allele among 2 or 3 outgroups. We then re-calculated the genetic load for a phylogenetically diverse group of species (the Przewalski’s horse, wolf, human and house mouse). Circles represent individual estimates and dotted lines depict the population averages for the different number of used outgroups. Although the population averages are higher when using only one outgroup, the differences are minimal and consistent across species.

**Fig. S9.**
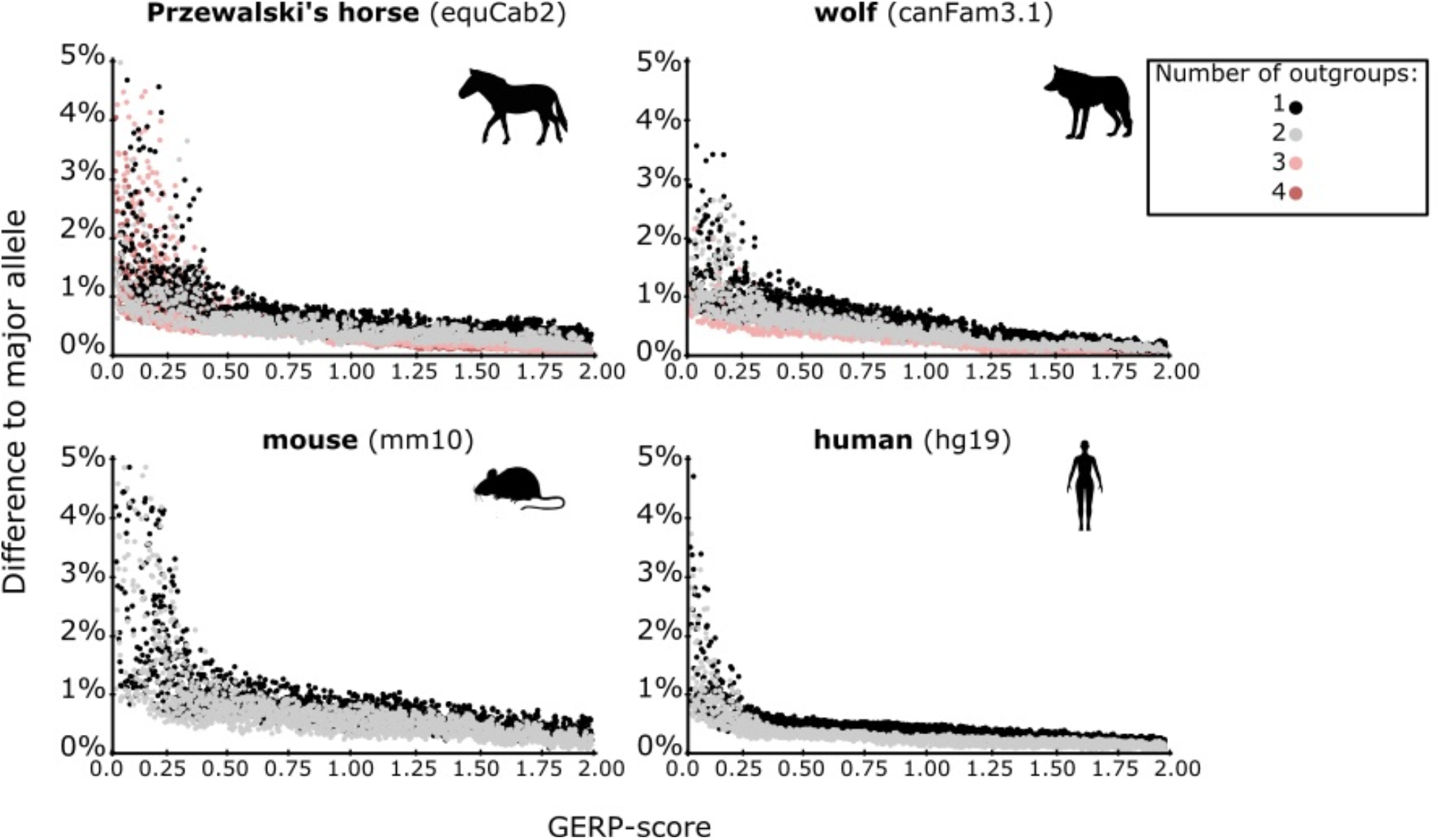
Ancestral allele inference depending on the number of used outgroups. Plots show the percentage of nucleotide differences to the major allele (among the complete phylogeny, e.g. all species that mapped to the site) by GERP-score depending on the number of outgroups used to infer the ancestral allele. Increasing the number of outgroups only slightly increases the likelihood of calling the correct ancestral allele and comes at the cost of having fewer sites in total.

**Fig. S10.**
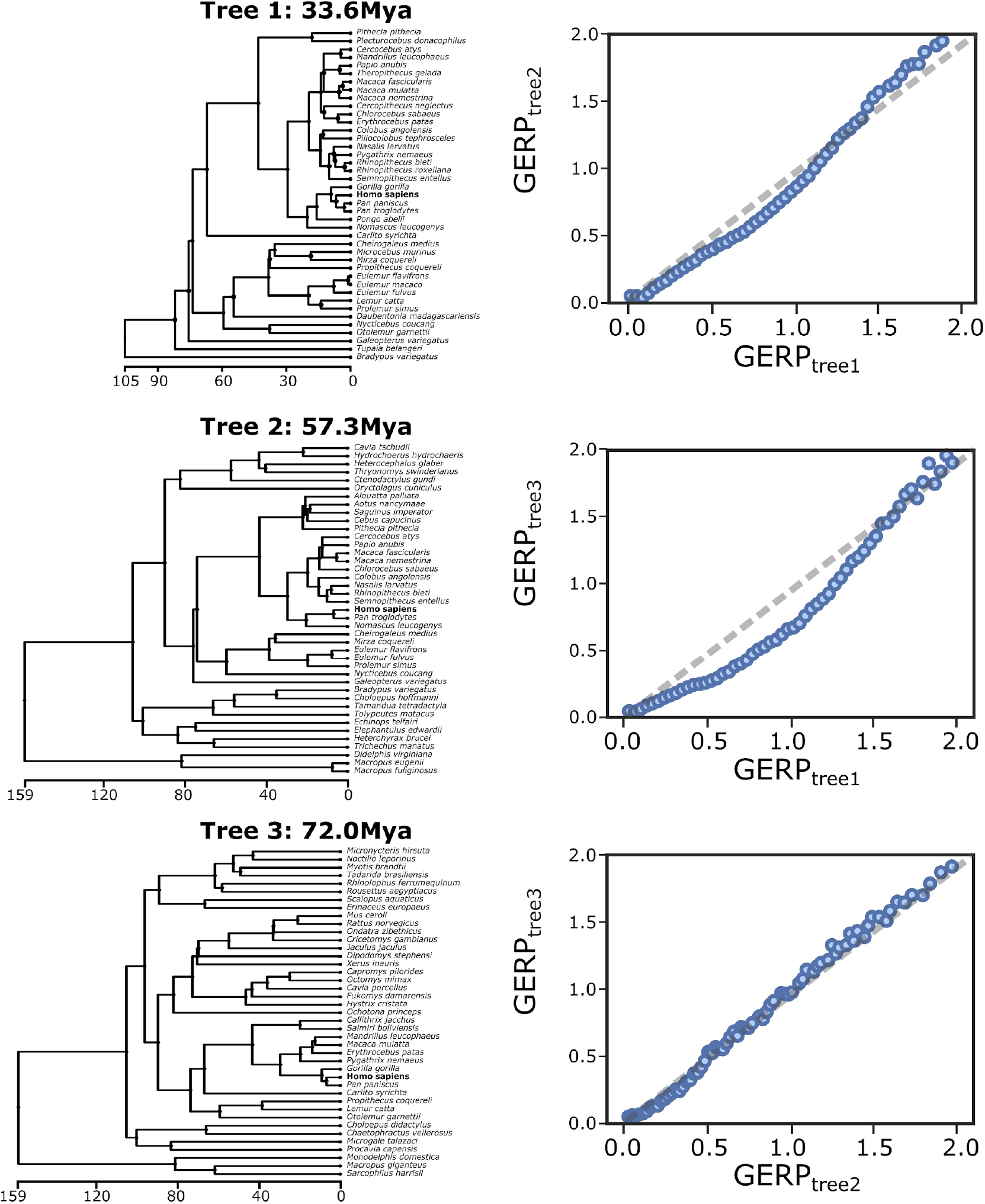
Correlation between GERP-scores calculated for the human genome (highlighted in bold) based on different species trees. The average divergence of the reference species to the human genome is depicted above each tree. The dotted grey line in the right-hand graphs depicts parity. GERP-scores remain similar despite the use of different reference species.

**Fig. S11.**
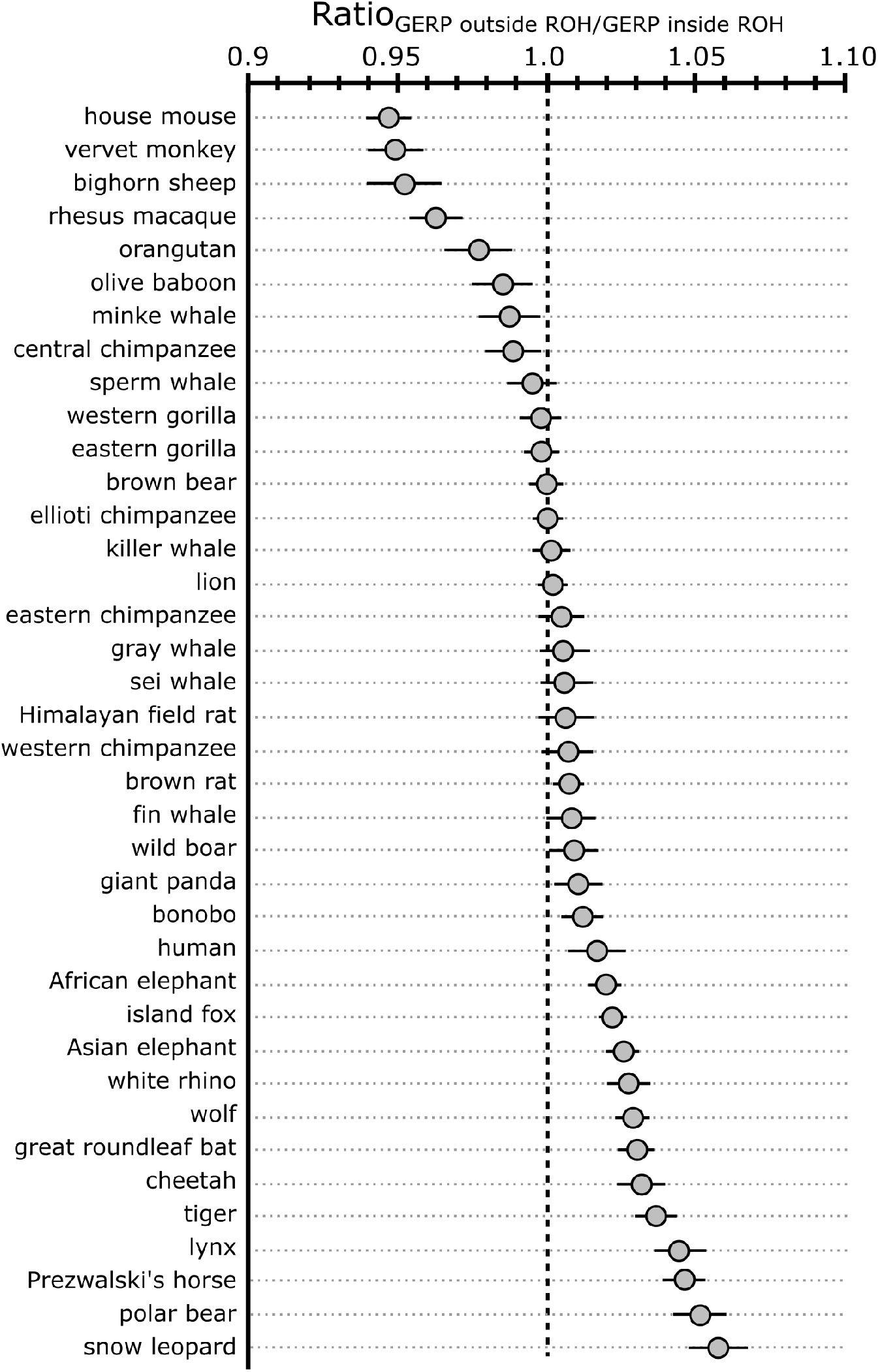
Ratio between estimates of genetic load within and outside of runs of homozygosity. A ratio below 1 corresponds to higher genetic load within the runs of homozygosity than outside of them, ratio above 1 corresponds to higher genetic load outside the runs of homozygosity compared to within. Bars show inter-individual variation, the position of the grey circle corresponds to the mean value per species. Species are ordered by increasing ratio. *\*Ratios were only calculated for species with high coverage genomes available in order to accurately infer ROHs*.

**Fig. S12.**
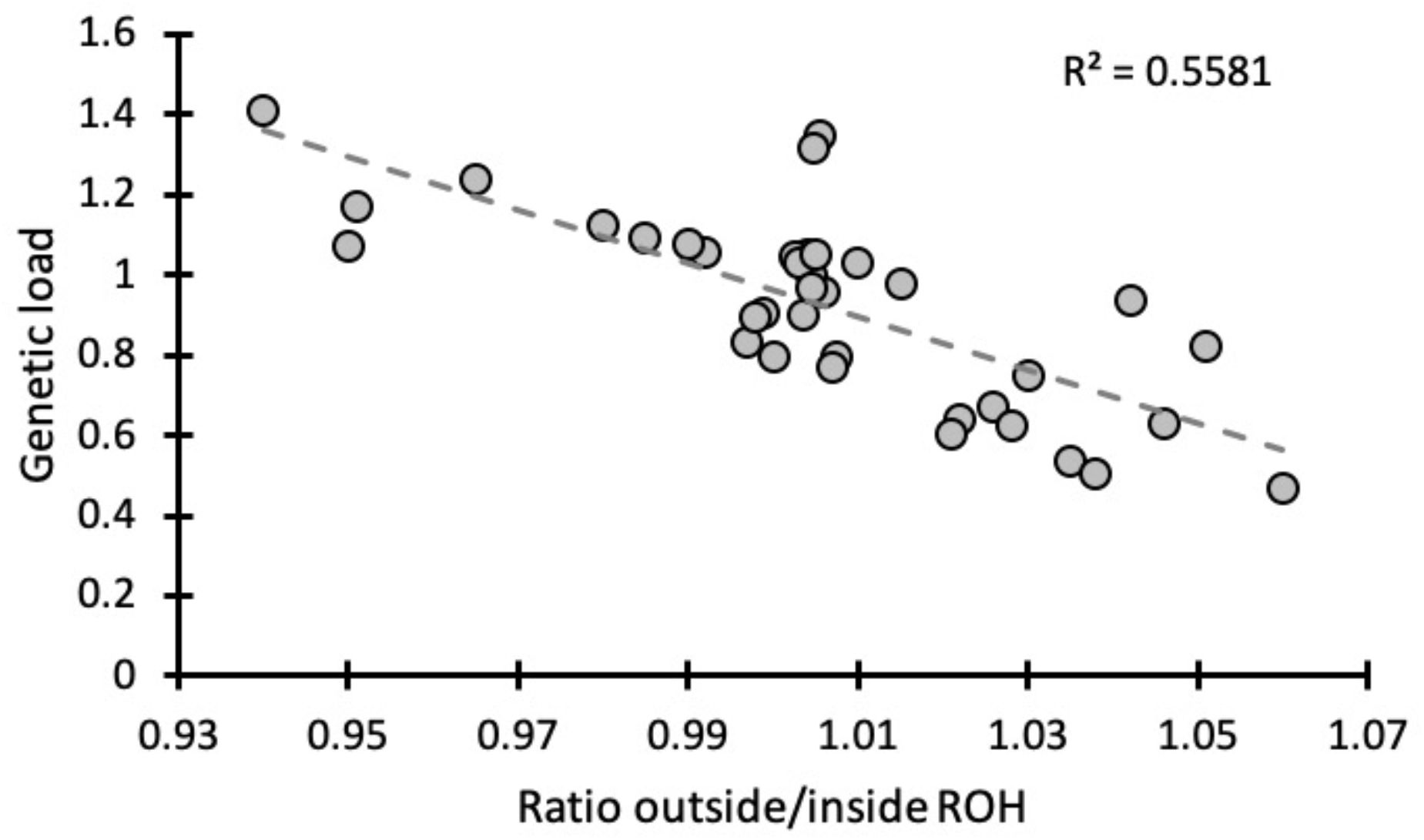
Correlation between overall genetic load and the ratio of load inside and outside ROHs. Each dot represents the species average. Species with lower load generally have fewer deleterious alleles inside ROHs than outside, whereas species with high genetic load carry relatively more deleterious alleles inside ROHs.

**Fig. S13.**
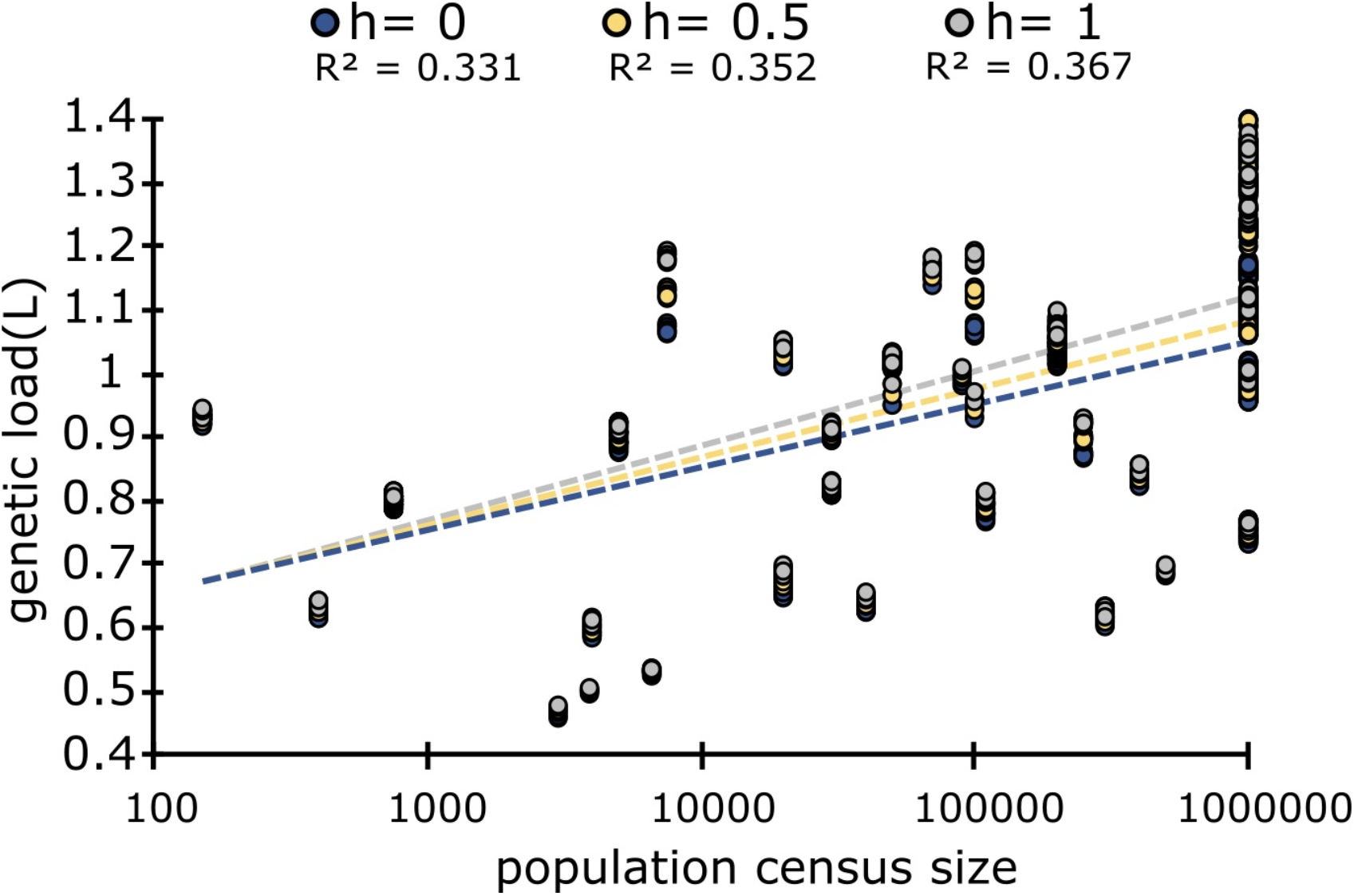
Relationship between genetic load and population size for different dominance coefficients. Each circle corresponds to a study population or species, the colour of the circle corresponds to the dominance coefficient. Genetic load was calculated with the formula L = 1-W = 2shx(1-x) + sx^2, following (1), where s=GERP score of the derived allele used as a proxy for the selection coefficient, x=allele frequency and h=dominance coefficient that ranges from 0 (recessive) to 0.5 (additive) to 1 (dominant). Genetic load increases with population size for all values of h, but the correlation is stronger for dominant alleles. For small populations, dominance coefficient has no effect on the estimates of genetic load, as most derived alleles are homozygous and thus exposed to selection independent of the dominance coefficient. In large populations the genetic load is higher if the derived alleles are dominant, but the effect of dominance is rather low in a cross-species comparison.

**Fig. S14.**
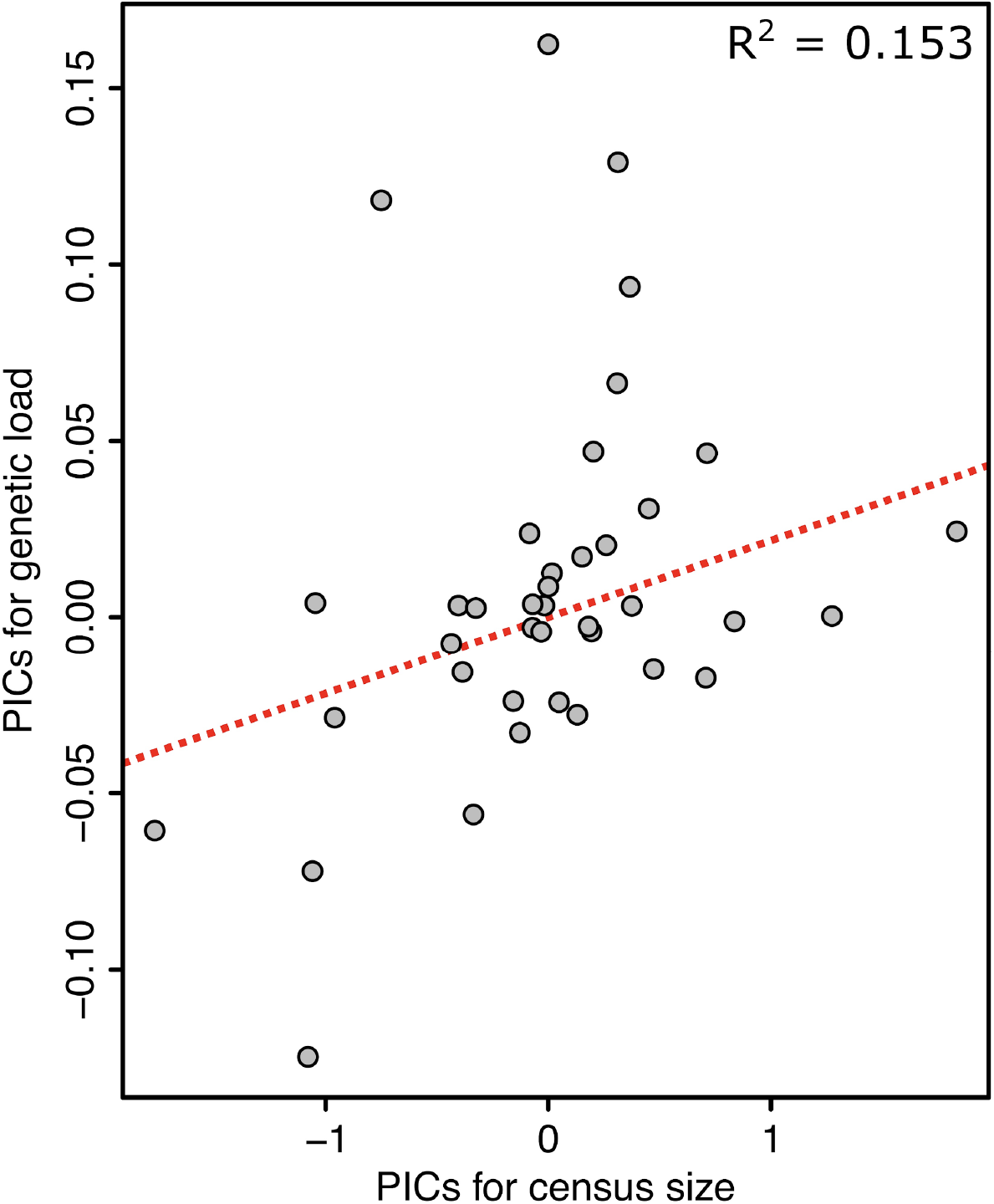
Phylogenetic independent contrast (PIC). Census size and genetic load are correlated, corrected for phylogenetic relationship between the species (2) to account for ancestral traits shared between closely related species.

**Table S1**.

**Individual genome re-sequencing data used to estimate genetic load and FROH**

(Provided as a separate file)

**Table S2**.

**Reference genomes used to calculate GERP-scores**

(Provided as a separate file)

## Notes

### Competing Interest Statement

The authors have declared no competing interest.

